# Ancient methicillin-resistant *Staphylococcus aureus*: expanding current knowledge using molecular epidemiological characterization of a Swiss legacy collection

**DOI:** 10.1101/2023.04.27.538514

**Authors:** Vanni Benvenga, Aline Cuénod, inithi Purushothaman, Gottfried Dasen, Maja Weisser, Stefano Bassetti, Tim Roloff, Martin Siegemund, Ulrich Heininger, Julia Bielicki, Marianne Wehrli, Paul Friderich, Reno Frei, Andreas Widmer, Kathrin Herzog, Hans Fankhauser, Oliver Nolte, Thomas Bodmer, Martin Risch, Olivier Dubuis, Sigrid Pranghofer, Romana Calligaris-Maibach, Susanne Graf, Sarah Tschudin-Sutter, Vincent Perreten, Helena M.B Seth-Smith, Adrian Egli

## Abstract

Few methicillin-resistant *Staphylococcus aureus* (MRSA) from the early years of its global emergence have been sequenced. Knowledge about evolutionary factors promoting the success of specific MRSA multi-locus sequence types (MLSTs) remains scarce. We aimed to characterize a legacy MRSA collection isolated from 1965 to 1987 and compare it against publicly available international and local genomes.

We accessed 451 ancient (1965-1987) Swiss MRSA isolates, stored in the Culture Collection of Switzerland. We determined phenotypic antimicrobial resistance (AMR) and performed Illumina short-read sequencing on all isolates and long-read sequencing on a selection with Oxford Nanopore Technology. For context, we included 103 publicly available international genomes from 1960 to 1992 and sequenced 1207 modern Swiss MRSA isolates from 2007 to 2022. We analyzed the core genome (cg)MLST and predicted SCC*mec* cassette types, AMR, and virulence genes.

Among the 451 ancient Swiss MRSA isolates, we found 17 sequence types (STs) of which 11 have been previously described. Two STs were novel combinations of known loci and six isolates carried previously unsubmitted MLST alleles, representing five new STs (ST7843, ST7844, ST7837, ST7839, and ST7842). Most isolates (83% 376/451) represented ST247-MRSA-I isolated in the 1960s, followed by ST7844 (6% 25/451), a novel single locus variant (SLV) of ST239. Analysis by cgMLST indicated that isolates belonging to ST7844-MRSA-III cluster within the diversity of ST239-MRSA-IIII. Early MRSA were predominantly from clonal complex (CC) 8. From 1980 to the end of the 20^th^ century we observed that CC22 and CC5 as well as CC8 were present, both locally and internationally.

The combined analysis of 1761 ancient and contemporary MRSA isolates across more than 50 years uncovered novel STs and allowed us a glimpse into the lineage flux between Swiss and international MRSA across time.

## Background

The introduction of penicillin during the 1930s and early 1940s was quickly followed by the rise of penicillin resistant *Staphylococcus aureus* due to penicillinase *blaZ* (1–3). Methicillin, a penicillinase-resistant antibiotic, was introduced in 1961, and within a year, the first methicillin resistant *S. aureus* (MRSA) isolate carrying the *mecA* gene emerged and established itself as a major nosocomial pathogen (4). The *mecA* gene is found on a mobile genetic element, the Staphylococcal Cassette Chromosome mec (SCC*mec*), of which different types have evolved, and which spreads via horizontal gene transfer (5). The acquisition of *mecA* has been dated back to the mid-1940s (6), and it is hypothesized that the SCC*mec* cassette originated from the *mecA* gene of *Staphylococcus fleurettii* and its surrounding chromosomal region (7). While one study advanced the hypothesis that the acquisition of the SCC*mec* cassette was a single evolutionary event (8), more recent studies supported the theory that MRSA emerged multiple times through independent events (9–11) and that cassette substitution can also occur, although far less frequently than cassette acquisition (12).

Molecular typing of MRSA isolates is a very important part of any local, national, and international epidemiology management strategy (13). Current unified MRSA clone nomenclature incorporates information on the genomic ancestry as multi-locus sequence typing (MLST), methicillin resistance status, and the SCC*mec* cassette type (e.g. ST250-MRSA-I or ST8-MSSA) (9).

The emergence of antibiotic-resistant *S. aureus* throughout the twentieth century can be roughly classified into four waves. First, the advent of penicillin resistance provided the first resistant *S. aureus* (1940-1960). Second came methicillin resistance, mostly linked to European hospitals (1960-1980), which spawned the term hospital-associated MRSA (HA-MRSA). In the third wave methicillin resistance spread worldwide (1980-1990). The fourth wave defines the rise of community-associated and livestock associated MRSA (CA-MRSA, LA-MRSA) (1990-2000) (14), coinciding with the development of resistance to ciprofloxacin and vancomycin (15,16). During this time, major clonal populations with specific STs emerged and disappeared through displacement by new, more successful epidemic clones (17). The first epidemic MRSA clone was identified as ST250 in the 1960s, within clonal complex (CC) 8. It is hypothesized that it arose as a single locus variant (SLV) of ST8-MSSA, which then acquired the SCC*mec* type I (9) and spread throughout Europe. ST250-MRSA-I was subsequently replaced by its SLV ST247-MRSA-I, first in Denmark around 1964 (18) and then globally, gaining resistance to further antimicrobials (9). ST247-MRSA-I was later displaced in Europe by other successful lineages in the late 1990s. ST239-MRSA-III, which also evolved from ST8-MRSA-III (9), was first identified in Australian hospitals in 1979 (19). Subsequently it was found in Brazilian hospitals in 1992 (20) and competed with ST247-MRSA-I in Portuguese hospitals throughout the 1990s (21,22). Later in the century, ST8-MSSA also acquired resistance to methicillin: one of the most successful contemporary CA-MRSA, USA300, belongs to ST8 (23,24). US300 has become endemic in North America and is still finding niches to colonize around the globe, with repeated introductions of USA300 into Europe reported (25–29). ST22-MRSA-IV, another global lineage, has been reported in many countries since the 1990s. ST22-MRSA-IV became a dominant HA-MRSA in England (UK-EMRSA-15) (30–32), and by the year 2000 was responsible for 65% of British MRSA bacteremia episodes (33). ST5 is a pandemic lineage reported in Europe, Asia, and North America (31). Previous comparative WGS studies have highlighted a recent increase in MRSA diversity, combined with success in ecological niches (30,34). This success is bolstered by new genetic elements associated with either increased invasiveness or lower fitness cost (26,35).

The molecular epidemiological situation of MRSA within Switzerland in a historic and international context has so far not been evaluated. The Culture Collection of Switzerland (CCoS)(36) is a national repository for microorganisms and contains ancient clinical *S. aureus* isolates collected between 1965 and 1987 based on unusual phenotypes, such as antibiotic resistance. This is the time-period when MRSA was establishing itself as a major nosocomial pathogen, from when the number of publicly available whole genomes is quite limited, with just 103 from isolates between 1960 and 1992. Therefore, we aimed to perform an in-depth characterization of this Swiss legacy collection using whole genome sequencing and to position the results in a global context through comparison with publicly available genomes and modern Swiss MRSA.

## Results and discussion

### Epidemiological context of the CCoS

We collected and analyzed a legacy MRSA collection from Switzerland (CCoS; n=451, collected between 1965-1987), all publicly available genomes from a similar time-period (public repositories; n=103, mainly from UK, Denmark, and Asia, collected between 1950 and 1992), and modern Swiss MRSA isolates collected at the University Hospital Basel (USB; n=1207, deriving from different Swiss institutions, collected between 2009 and 2022). We characterized the genomes using MLST, core genome MLST (cgMLST), and whole genome phylogeny.

The complete dataset of 1761 genomes comprise 81 STs, of which 75 have been previously described (**Figure 1**, **Supplementary material 1**). Prior to 2000, 554 genomes were analysed, belonging to 17 STs. Of these, six STs were previously undescribed and include 32 isolates. These six undescribed STs comprise two novel combinations of known loci (ST7844 and ST7842) and four exhibiting previously undescribed alleles (ST7638, ST7837, ST7839, and ST7843) **(Figure 1).**

**Figure 1:**
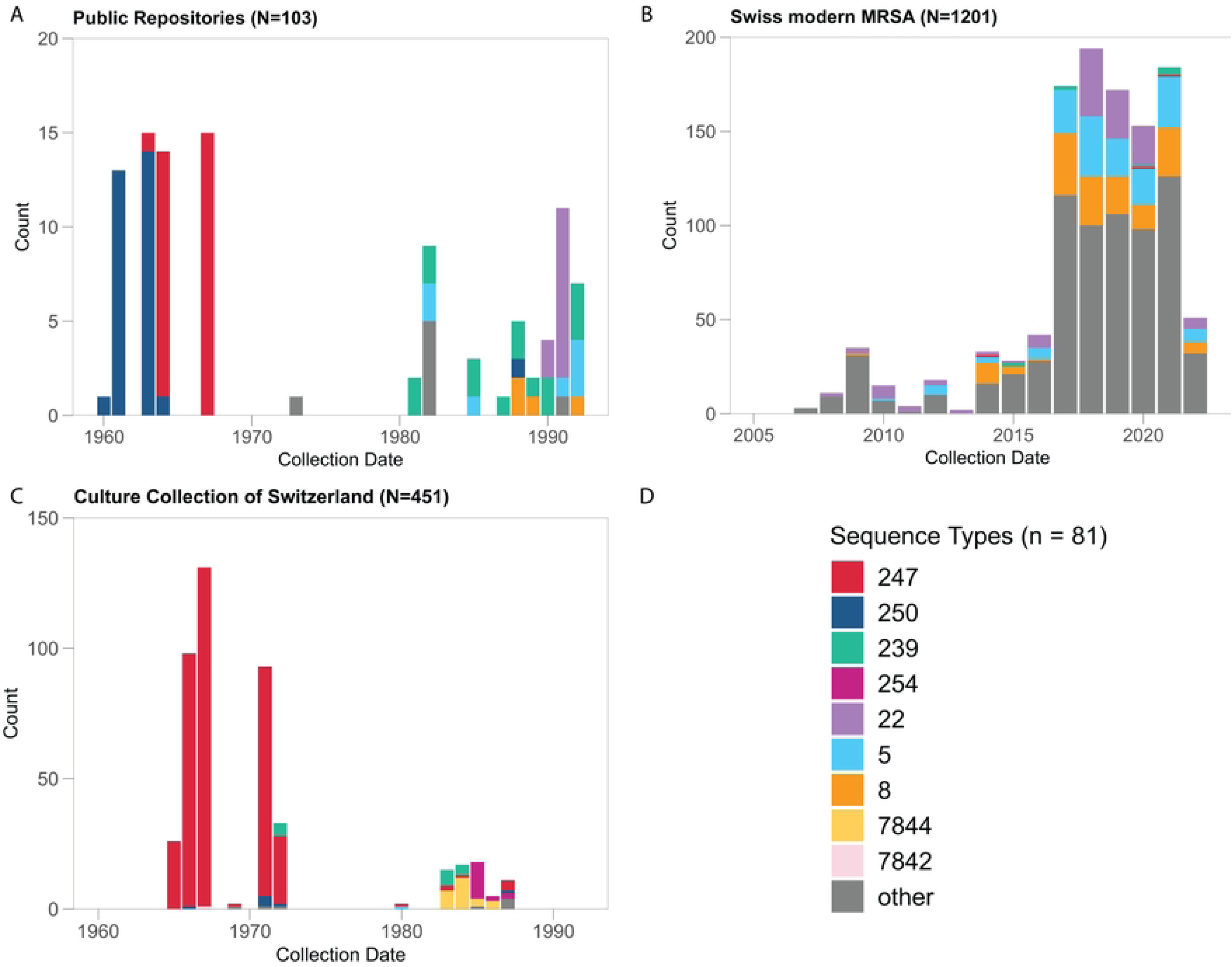
Epidemic distribution of MLST sequence types by collection year and dataset (A: Public repositories, B: Culture Collection of Switzerland, C: Contemporary Swiss MRSA, D: Legend). Samples with no collection year are not represented, n=6. STs present in low numbers are only shown as “other” (gray). Be aware that each subfigure has a different y-axis.

In both ancient datasets (**Figure 1A, 1C**) most of the samples were collected between 1960 and 1972. The CCoS collection was dominated by ST247 isolates from the mid 1960s to the early 1970s, coinciding with low levels of ST250 and ST239 (**Figure 1C**). A decade later, these STs were still present but we also registered ST254 and ST7844. The STs from public repositories of the same time period are similar (**Figure 1A**). From 1980 onwards, the STs represented in the international collection are ST239, ST8, ST5, and ST22. Of note, all the previously mentioned STs, except for ST5 and ST22, belong to CC8, as seen in the MLST minimum spanning tree (MST, **Figure 2**). Among modern Swiss MRSAs, we observed many more STs with the most common being ST22, ST5, and ST8 (**Figure 1B**). Other STs common in the CCoS are detected in small numbers among modern samples (ST239 and ST247).

**Figure 2:**
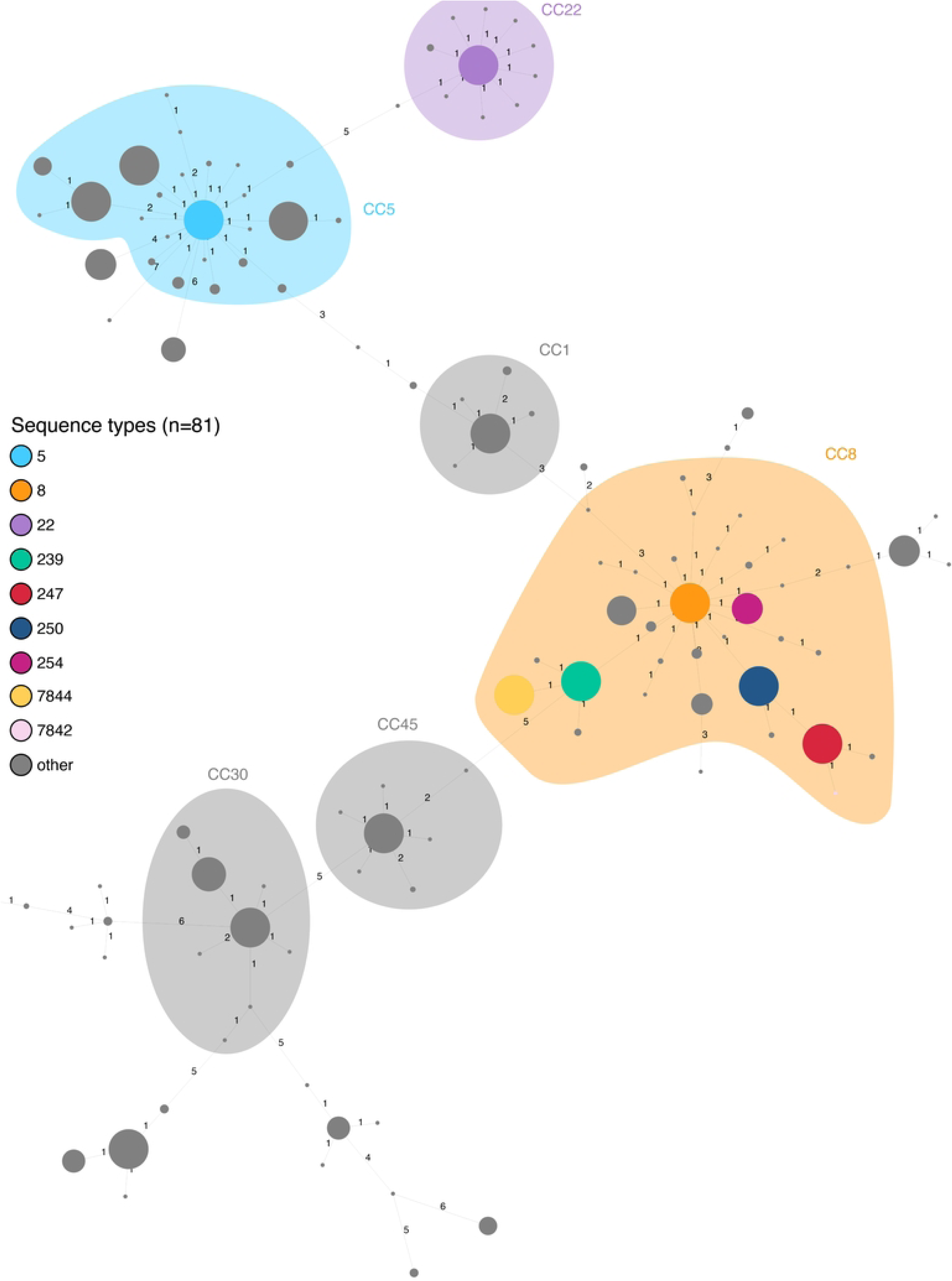
MLST minimum spanning tree generated by Ridom SeqSphere+ (n=1761). Nodes colored by ST with CC superimposed. CCs are defined as all STs which match their central genotype (ST) in four or more loci. Branch labels show number of allelic differences.

Within CC8, ST250, ST254, and ST239 are single locus variants (SLVs) of ST8, while ST247 is a SLV of ST250. The novel STs 7844 and 7842 are SLVs of ST239 and ST247, respectively. The high prevalence of ST247 among the CCoS sample set mirrors the epidemiological situation of other European countries at the time (9,18,37). In addition, the Swiss ST239 from 1972 (n=5) demonstrate that ST239 isolates were already present in Europe in the early 1970s and provides the earliest sequenced ST239 MRSA available (19). The results were visualized on a map **(Supplementary material 14).** An interactive version of the map can also be found online under https://github.com/svannib/ancient_MRSA/tree/main/ancient_MRSA_map.

### Identification of successful ancient lineages through cgMLST clustering

The average *S. aureus* genome contains 2872 coding sequences (CDSs), of which 1861 are present in the *S. aureus* core genome MLST scheme (38,39). The cgMLST analysis of all ancient genomes (n=554) yielded a cgMLST MST whose nodes have been colored by ST (**Figure 3**), isolation country (**Supplementary material 2**), and collection decade (**Supplementary material 3**). Isolates of the same ST can be seen to generally cluster together. However, we observed that some STs exhibit higher diversity between isolates than others. For example, distances among ST247 isolates rarely exceed 30 alleles, while ST5 samples often lie more than 200 alleles apart. The isolates of these STs were sampled over very different time periods (ST247 mainly from 1963 to 1987 with three sample after 2000, while ST5 is more evenly distributed from 1980 to 2022, both with a gap in from mid-1990s to the mid-2000s) so this wide sampling could explain, at least in part, this higher diversity. Generally, STs with higher diversity contain samples isolated over a longer period. Another example is ST239, which shows a high diversity by cgMLST (allelic divergence between 50 and 120), with isolates from geographically distant countries (Australia, Singapore, United States, and Switzerland) and collected between 1972 and 2021. The genomes from the earliest ST239 (n=5) are central to this group (**Supplementary figure 2**, highlighted in red). ST5 isolates are distant from other STs (1382 allele differences), displaying allelic distances up to 207 within this ST. ST22 samples are also distant from other groups (1599 allele differences), were isolated from different regions of Britain, and show allele differences of up to 37. The biggest allelic differences are seen between CCs, exceeding 1000 allele differences in all cases, while inside CCs distances never exceed 500 allelic differences.

**Figure 3:**
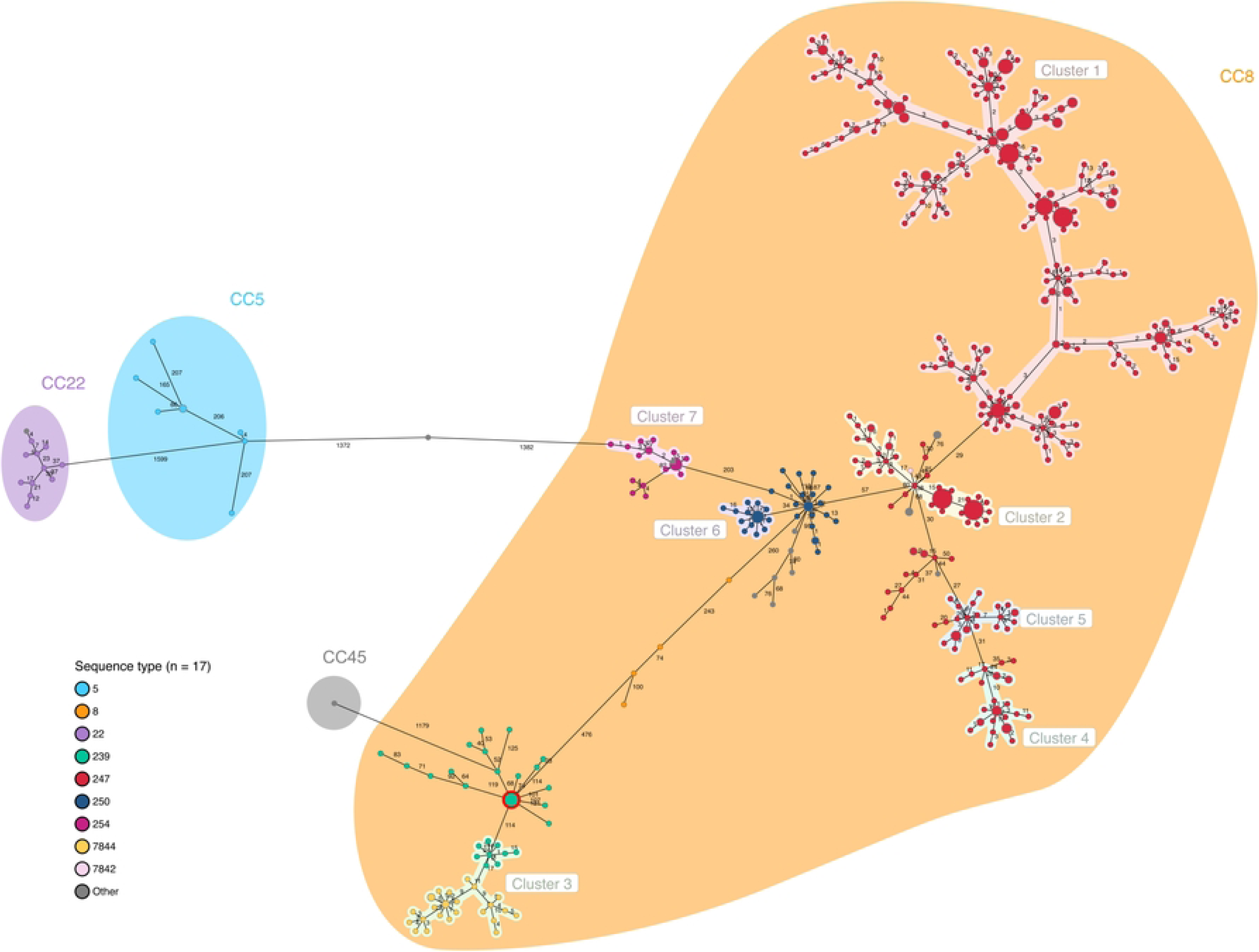
cgMLST MST of CCoS and public repository genomes of MRSA (1960-1992, n = 554), nodes colored by sequence type and CCs are shaded. Clusters are shown with a maximal cluster distance of 24 allele differences and a minimal cluster size of 15. The earliest ST239 (1972) isolates are circled in red.

We identified seven putative transmission clusters, using a cluster cutoff of 24 allele differences (**Figure 3**). Cluster 1 contains ST247 samples collected between 1960 and 1999, suggesting that isolates within this cluster remained successfully circulating throughout the decades. Cluster 2 also encompasses ST247 samples isolated in the 1960s from Switzerland and Denmark, where the first ST247 was identified. Cluster 3 includes both Swiss (n=20) and British (n=1) ST239 isolates, along with ST7844, the two STs separated by 11 allelic differences. Clusters 4 and 5 contain Swiss isolates mainly from the 1960s and 1970s, respectively. Cluster 6 contains British ST250 samples from the 1960s, representing the first described MRSA. Cluster 7 covers German ST254 whose closest relative is ST250 (203 allelic differences), this ST is usually associated with infection in horses, and is rarely seen in humans (40).

### Changing epidemiological landscape at the turn of the century

Analysing all MRSA in our collection, from 1960 to 2022 (**Supplementary material 4**), we observe that CC8, dominant in 1960s-1970s, is not the main CC represented across modern samples. The only ST within CC8 which plays a main role after the turn of the century is ST8 (n=141/1207, 12% of modern Swiss samples). This might be due to the international predominance of ST8 CA-MRSA in the last decades (41). Particularly interesting is the strong modern presence of CC5 (n=306/1207, 25% of modern Swiss samples) and CC22 (n=210/1207, 17% of modern Swiss samples) isolates, which were present at the international (but not Swiss) level in the 1980s and 1990s. This suggests that international lineages arrived in Switzerland around the turn of the century. Furthermore, other complexes are present in modern Swiss hospitals such as CC1 (n=85/1207, 7%), CC30 (n=86/1207, 7%) and CC45 (n=53/1207, 4%).

In general, clones associated with HA-MRSA infections are the most prevalent both in ancient Swiss and public assemblies, possibly due to sampling bias. Modern global epidemiology shows less homogeneity, with lineages associated with community infections being more prevalent (42). These changes in epidemiological landscape mirror the recent growth in MRSA biodiversity reported internationally (30,34).

### Genotypic antimicrobial resistance (AMR) prediction and core genome phylogeny

To analyze the genomic data on a SNP rather than allele basis, core genome SNP phylogenies were calculated, calculated first on genomes from the CCoS and from public repositories sampled between 1960 and 1992 (n=554; 2034 core genes present in over 99% of 554 isolates) (**Supplementary material 5**), and subsequently with all MRSA covering a period from 1960 to 2022 (n=1761; 1225 core genes across 777 genomes) (**Supplementary material 6**). Both phylogenies show isolates from the same ST clustering together, as seen with cgMLST. We also observe the previously mentioned increased diversity among modern MRSA, with samples isolated after 2009 having a broader variety of STs.

Prediction of AMR from the genome was limited to antibiotics that are important: (i) for classification of MRSAs (oxacillin); (ii) for the history of MRSA (penicillin); or (iii) current clinically relevant antimicrobials (tetracycline, erythromycin, clindamycin, gentamicin, and ciprofloxacin). Sensitivity of the methods, the ability to correctly predict AMR, and specificity, the ability to correctly predict susceptibility, were calculated by comparing genotypic predictions to available phenotypic data (**Table 1**).

**Table 1:**
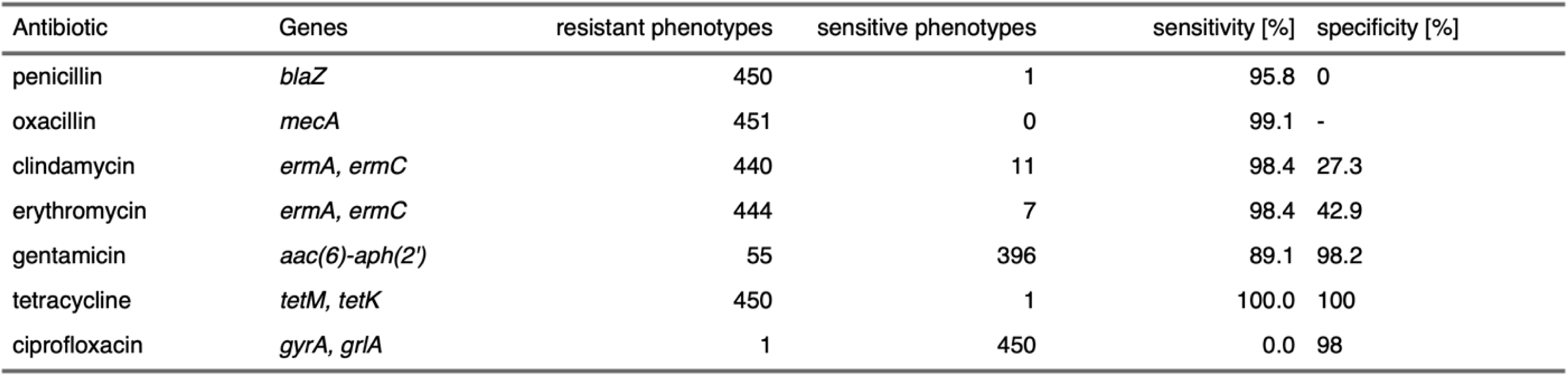
Sensitivity and specificity of AMR prediction for selected antibiotics and their respective antibiotic resistance encoding genes (ARG). The number of resistant and sensitive phenotypes given are based on the antibiograms (n = 451, EUCAST).

The detection of antibiotic resistance genes (ARGs) of *S. aureus* to predict phenotypic resistance can reach specificity and sensitivity similar to routine susceptibility testing (43,44). In our case, sensitivity to clindamycin and erythromycin seem to be particularly challenging to predict. Both phenotypic and predicted antibiotic susceptibility data are displayed adjacent to the core genome phylogeny in **Figure 4** with a binary heatmap showing concordance between the two to facilitate interpretation. As expected in an MRSA-only dataset, all isolates are resistant to meropenem, which was consequently removed from the visualization. However, four samples from the CCoS display an oxacillin-resistant phenotype despite no *mecA* or other *mec* variants being identified within the genomes. This could be due to beta-lactam resistance caused by mechanisms other than *mecA*, such as expression of penicillin-binding-proteins with low antibiotic affinity (45,46) or overexpression beta-lactamases (47) or loss of the cassette after the phenotype was determined.

**Figure 4.**
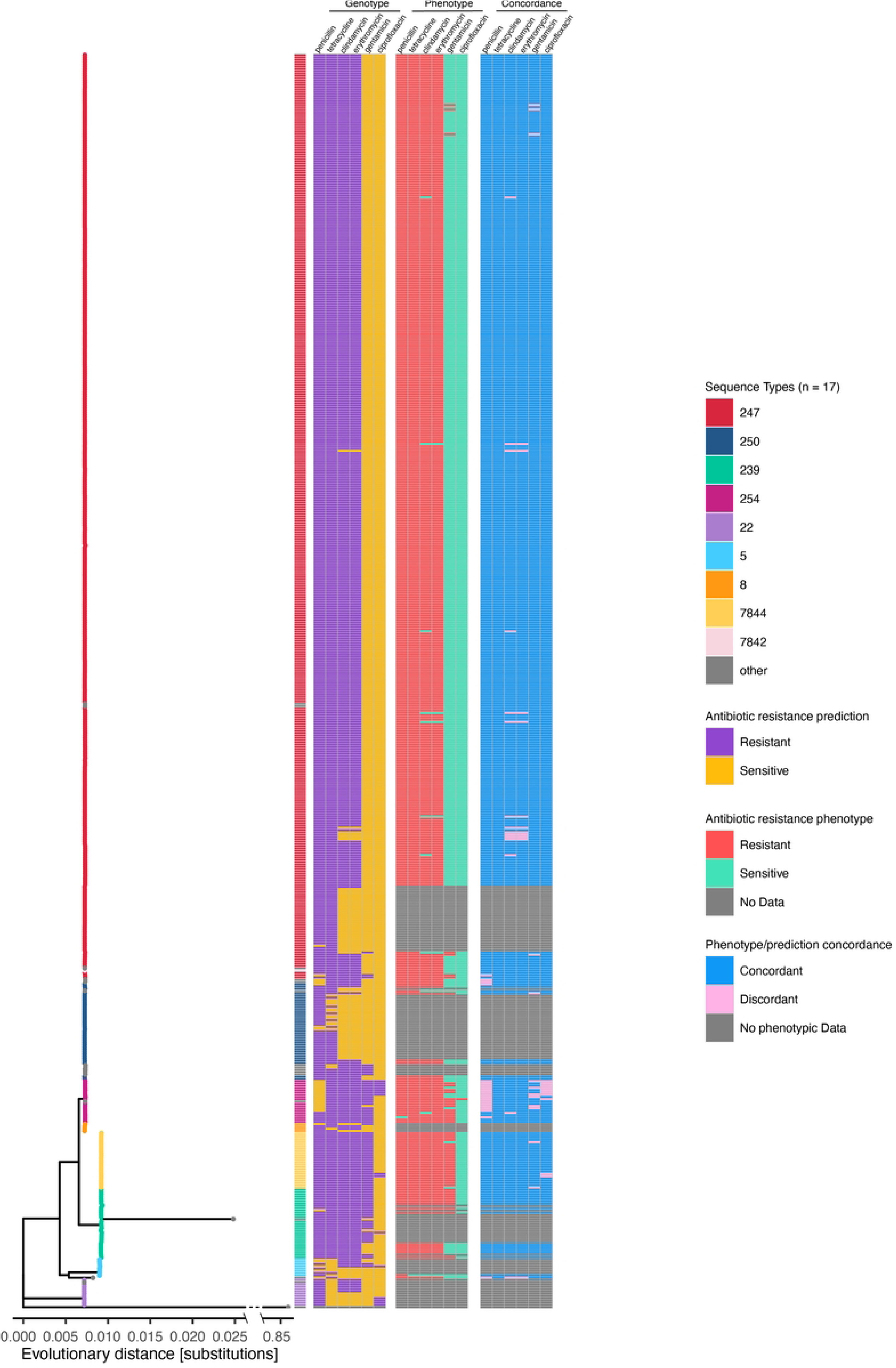
Predicted resistance/sensitivity, phenotypical resistance/sensitivity and concordance between the two mapped to a Maximum likelihood core genome SNP tree of ancient Swiss and international MRSA (1960-1992, n = 554). Leaves colored by sequence type. Outgroup (*S. epidermis*) line shortened through X-axis break for visualization purposes.

Penicillin resistance (*blaZ)* is widespread in this dataset (99.8%, 450/451 penicillin resistant phenotypes, 94.9%, 526/554 ancient genomes possess *blaZ*), being absent only from the majority of ST5 and some ST254 isolates (**Figure 4**). MRSA possessing *blaZ* has been previously documented: since the late 1960s to this day, isolates are often resistant to penicillin (48,49). Also, among modern Swiss MRSA, *blaZ* is still present in the majority of isolates (89.5%, 1080/1207 genomes), a finding supported by other studies on European *S. aureus* (50). Resistance to tetracycline is widespread (94.0%, 521/554) except for within ST5, ST22, and some ST250 genomes. Clindamycin resistance and erythromycin resistance are common among ST247 samples, except for ancient Danish genomes, present in some ST5 and ST22, and rare in ST250. Gentamicin resistance is a common feature among ST239, ST254, and ST7844, while being sparsely present throughout the rest of the dataset; ST22 and ST250 are uniformly gentamicin sensitive. Ciprofloxacin resistance is rare across the entire dataset. However, prediction of ciprofloxacin resistance is not very sensitive (**Table 1**), and the more recent introduction of the antibiotic on the market (1987) (51) might explain the limited numbers of resistant isolates in ST239, ST5, and ST22. Higher incidence of ciprofloxacin resistance is registered in modern Swiss MRSA (37%, 452/1207).

This dataset suggests that ancient ST247 gained genes leading to a broader resistance profile (clindamycin, erythromycin, and tetracycline) compared to some early ST250. This might be one of the factors which contributed to the success of the former and to the decline of the latter ST as epidemic MRSA clones (9). This is supported by other studies conducted on early MRSA isolates (6). The early Danish ST247 samples in this dataset are erythromycin and clindamycin sensitive, but they may present a biased portrayal of isolates of the time, since widespread erythromycin and clindamycin resistance in ST247 isolates from the same period have been reported (18). The phylogeny of the dereplicated dataset (**Supplementary material 7**) suggests that modern Swiss MRSA have a higher variability of resistance patterns than ancient MRSA, even among closely related isolates. This could be due to the isolates coming from geographically separated Swiss hospitals, or may be an artefact of the analytical dereplication performed on the modern Swiss MRSAs, as groups of very similar genomes are collapsed into one datapoint.

### Virulence genes

Across all datasets, 35 unique virulence encoding genes were identified, the most common being for hemolysins and proteases, present in more than 90% of all isolates. Also common were staphylokinase (*sak*), staphylococcal complement inhibitor (*scn*), and toxin encoding genes such as *lukD, lukE, sek, seq,* and *seb*. Other staphylococcal enterotoxins (*se_*) were far less common, while the arginine catabolic mobile element (*ACME*) and toxic shock syndrome toxin (*tst*) were present in under 1% of samples. A virulence gene presence/absence heatmap was mapped to a phylogenetic core genome tree of the isolates (**Figure 5**), showing frequent co-occurrence of *sak* and *scn*. Furthermore, international ST5 and ST22 isolates exhibit a general lack of serine-like proteases (*splA/B/E*) and toxins (*lukD, lukE, seb, sek, seq*) in favor of another group of staphylococcal enterotoxins (*seg, sei, sem, sen, seo, seu, sec, sel*). This strong co-occurrence to one another may suggest a pathogenicity island. ST239 and ST7844 broadly lack *seb* while generally displaying *sea* presence. However, ST7844 and its most closely related Swiss ST239 group also lack *sek* and *seq*. Previously described pathogenicity islands which could play a role in the distribution and dissemination of the virulence genes seen in the dataset are vSAβ (*sea, seg, sei, sem, sen, seo, splA, slpB, slpE, lukD, lukE*), saPI3 (*seb,sek,seq*), and plB485 (*sej, sed*) (52). The major toxin Panton-Valentine Leukocidin (*lukF-PV, lukS-PV*) which heightens the virulence of MRSA (53) is sparsely present in modern Swiss MRSA, but completely absent from isolates prior to 2009 (**supplementary material 8**). This lies in contrast with the high rates of Leukocidin reported in the US (54). A limitation of this virulome analysis is its reliance on WGS and gene presence/absence instead of diagnostic tests. Still, this approach has shown high concordance with phenotype-based methods (55).

**Figure 5:**
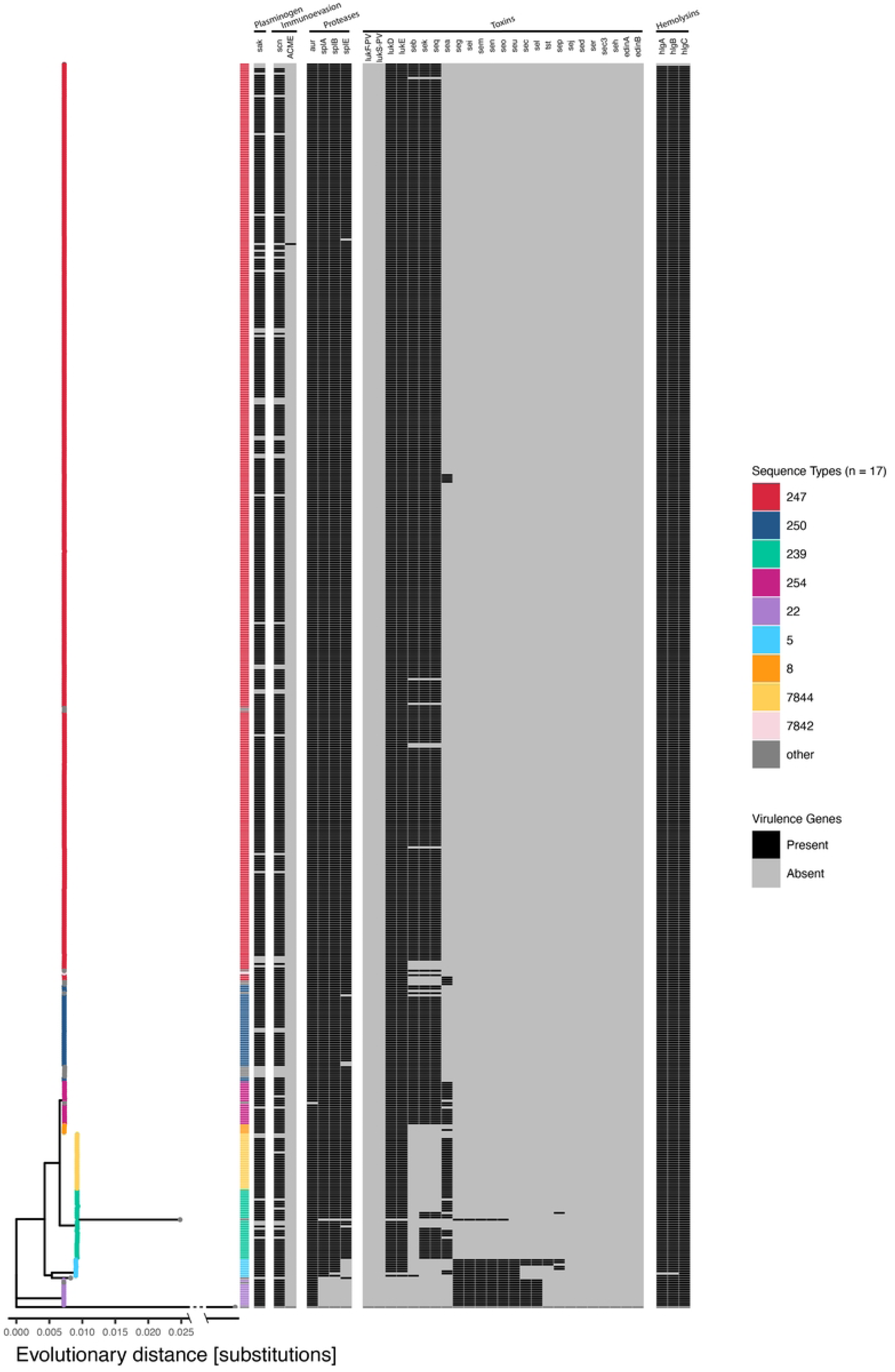
Virulence gene presence/absence heatmap mapped to a maximum likelihood core genes phylogenetic tree of ancient Swiss and international MRSA (n = 554). Leaves colored by sequence type. Outgroup (*S. epidermis*) line shortened through X-axis break for visualization purposes.

### SCC*mec* Types

Among 777 dereplicated genomes, six unique SCC*mec* cassette types were detected, as well as several ambiguous predictions. For isolates which were predicted as methicillin-sensitive despite having a resistant phenotype (n=5), no cassette was found in their genomes. SCC*mec* type I is by far the most prevalent (462/777, 59%), followed by type IV (144/777, 19%), and type III (61/777, 8%). SCC*mec* cassette types mapped to the core genome phylogeny (**Supplementary material 9**) illustrate strong co-occurrence between cassette types and ST lineages. SCCmec I is strongly represented within ST247 and ST250 among ancient isolates (27). SCC*mec* type IV (2B) is common among ancient and modern ST22, ST5, and ST8 isolates (**Supplementary material 10**). Three of the five major ST8-related epidemic MRSA clones are represented in this dataset (27). ST247-MRSA-I, ST250-MRSA-I (accounting for 80% (442/554) of the ancient genomes), and ST8-MRSA-IV. Another strongly represented epidemic MRSA, although it rose to prominence later than ST250-MRSA-I and ST247-MRSA-I, is ST239-MRSA-III (27), which has the same cassette as the SLV ST7844-MRSA-III.

In modern Swiss MRSA, *SCCmec* type IV in ST5, ST8, and ST22 isolates is the dominant cassette type. These lineages have often been reported as dominant in many countries spanning the globe (56–61). The wide presence of type IV in different MRSA lineages and other staphylococcal species might suggest an improved horizontal genetic transfer rate and/or lower fitness cost (35,62,63).

International lineages shaped modern Swiss MRSA epidemiology

Based on cgMLST and core SNP trees, three groups were chosen for whole genome SNP analysis: ST239, ST7844 and closest STs (group 1, **Supplementary material 11A-B**); ST250 together with ST247 (group 2, **Supplementary material 12A-B**); and ST22 (group 3, **Supplementary material 13A-B**). SNP phylogenies were generated using hybrid assembled Illumina/Nanopore reference genomes from within the cluster to best capture the diversity, and time trees were used to estimate the temporal origins of each group.

The resulting phylogenic tree of ST239 indicates that group 1 strains evolved from an ST239 ancestor, with SNP in MLST target genes resulting in SLV within the group. One example is a mutation in *glpf* which gave rise to ST7844 in a lineage which appears to previously have been successful, but which is not represented in modern Swiss MRSA.

One cluster within this group contains Swiss, German, and British ST239 and ST7844, which were collected before 1990 (Bayesian cluster 3). It has no modern samples, hinting at its disappearance from Switzerland. Another cluster contains the earliest ST239 from Switzerland and international isolates from the 1980s and 1990s (cluster 4). Interestingly, some modern ST239 isolates (cluster 1-2) are closer to cluster 4 than to other modern samples. Cluster 5 covers both ancient ST239 from Singapore and modern Swiss ST368 and ST241, pointing to an introduction of this group into Switzerland at the turn of the century. The root of group 1 was dated to approximately 1950 (95% confidence interval (CI): 1941-1957).

The SNP phylogeny of ST247 and ST250 (group 2) shows that six Bayesian clusters of ST247 are present in the CCoS (Supplementary File 12A-B). Bayesian cluster 6 covers both Swiss and Danish samples collected in the 1960s, alluding to international transmission within Europe at the time. Modern ST247 play a small role in the modern Swiss epidemiology of MRSA, but these modern isolates cluster with ancient Swiss, British, and Belgian ST250s. The root of this group was dated to 1870 (95% CI: 1851-1885).

The ST22 phylogeny (group 3, Supplementary material 13A-B) (origin dated to 1914 (95% CI: 1907-1918)) again hints at the international origin of lineages circulating today in Switzerland, as British isolates from the 1980s are closely related to the ancestors of 85 modern Swiss samples (clusters 5 and 4).

Overall, SNP phylogeny of cgMLST clusters provides evidence as to which of the old international MRSA lineages appeared briefly in Switzerland before being displaced, and which contributed to the MRSA diversity we see in modern Switzerland.

## Conclusion

Since the 1960s, MRSA has been a challenging bacterial pathogen faced by clinicians worldwide. This study sheds light on the spread and relationships of major early MRSA clones. Our genome collection includes 451 MRSA samples from CCoS isolates between 1965 and 1987s, alongside 103 ancient MRSA genomes from public repositories and 1207 modern MRSA isolated in Swiss hospitals. Despite being a sample set which is potentially unrepresentative of the diversity in Switzerland at the time, our data reveal an ancient epidemiological landscape within Switzerland similar to that in the rest of Europe at the time. MRSA lineages which played an important role across European and Swiss hospitals from the 1960s to the 1990s, such as ST247-MRSA-I, ST250-MRSA-I and the earliest ST239-MRSA-III are represented in the CCoS. Today, these clones appear to have been displaced in Switzerland, with international lineages from the last quarter of the 20th century, including ST5-MRSA-IV, ST8-MRSA-IV, and ST22-MRSA-IV now the major players in Swiss hospitals. An analysis of the AMR and virulence profiles showed how different STs are associated with different AMR and virulence encoding genes. The limitations of this study are: a) the lack of sequenced Swiss isolates between 1988 and 2008, which prevents us from fully understanding the epidemiological changes which happened over the turn of the century and b) the lack of phenotypic resistance data for the current Swiss MRSA, which would make our lineage-resistance association more robust and precise. These data from CCoS hold high scientific interest, as the collection contains some of the first MRSA ever isolated and whose whole genome sequences we now present. We have thus significantly increased the public available genomes from the early period. The volume of isolates and phenotypic characterization make them an important addition to the pool of MRSA genomic data of isolates isolated in the second half of the 20^th^ century. The analysis of early pathogenic MRSA such as these isolates leads to a deeper understanding of its epidemiology which may help current and future efforts in infection control and prevention.

## Materials & Methods

### Whole genome sequencing and phenotypic AMR profiling

Bacterial genome assemblies from the CCoS and from the University Hospital Basel (USB) were generated at the Division of Clinical Bacteriology (USB) according to their ISO 17025 accredited standard procedures. DNA was extracted with a Qiagen BioRobot EZ1 using the QIAamp DNA Mini Kit (QIAGEN, Hilden, Germany) and according to the manufacturer’s guidelines. Library preparation was performed using an Illumina Nextera DNA Flex Library Prep Kit and multiplexing at 96-plex on a NextSeq 500 System using the Mid-Output Kit (Illumina, San Diego, USA). Resulting fastq files underwent a quality check, where the sequences with a phred score lower than 30 were discarded (only sequences with a base calling accuracy above 99.9% are kept). Sequencing data was quality controlled using FastQC (v 0.11.5) (64), MetaPhlAn (v 2.7.7) (65). Adaptors were trimmed using Trimmomatic (v 0.38) using default parameters (ILLUMINACLIP:2:30:10 SLIDINGWINDOW:4:15 MINLEN:125) (73). Genome assemblies were created *de novo* using Unicycler (v 0.3.0b) (66) and checked with QUAST (v 5.0.2) (67) Further details can be found in (68).

For the isolates from CCoS, a phenotypic antibiotic resistance analysis was conducted as follows: bacteria were grown on CHROMID/MRSA Agar (bioMérieux, Marcy-l’Étoile, France) at 35 ± 2°C for 48h. Antibiotic susceptibilities were determined using a microdilution assay (Vitek2, bioMérieux, Marcy-l’Étoile, France) and interpreted according to breakpoints published by the European Committee on Antimicrobial Susceptibility Testing (EUCAST), version 9.0 (69).

### Nanopore sequencing

Ten isolates from the CCoS were chosen for nanopore sequencing. DNA was extracted from the bacterial pellet using the DNA mini (Qiagen) kit on the QIAcube robot (Gram + Enzymatic lysis protocol) with 150 μl elution volume. Long read libraries were prepared using the ligation sequencing kit (SQK-LSK 109). The libraries were sequenced using GridION with R9.4 flow cells with a default 72h run time. Basecalling and de-multiplexing were carried out within the inbuilt MinKnow (21.05.12) software. Basecalling was done using Guppy (5.0.12) high accuracy model. Quality check was carried out using Nanoplot (1.35.4) (70) and MetaPhlAn (3.0.13) (65). Trimming was done with Porechop (0.2.3) (71). Short reads were removed with Filtlong (0.2.0) (72). Assembly was performed with flye (2.8.1) (73) and polished with racon (1.4.7) (74), medaka (1.4.4) (75) and polypolish (v0.4.3) (76). Raw data can be found at the European Nucleotide Archive under accession number PRJEB59014.

### Collection of public genomes

The timeframe covered by the 451 *S. aureus* isolates from the CCoS is from 1965 to 1987. The parameters for the search of public genomes were set at a slightly wider timeframe (1960-1992) to allow for a wider selection of isolates. 108 assembled genomes were downloaded from NCBI Pathogen Detection (77). NCBI Pathogen. The 108 genomes, and their respective metadata, were obtained by applying the following criteria: Species: *Staphylococcus aureus*, Collection date: 1960 to 1992. 150 genomes were downloaded from Pathogenwatch (78,79). The assemblies were found by searching for *Staphylococcus aureus* genomes collected between January 1960 and December 1992.

### Data filtering

Filtering criteria were applied to both the public and in-house sequenced isolates. Only isolates which satisfy the following conditions were kept: Genome length within ± 10% of reference *S. aureus* genome NCTC 8325; where recorded, read depth greater than 20x; if antibiogram was present, resistance to oxacillin, otherwise presence of *mecA* in the genomes was considered as denoting an MRSA. 46 isolates with an oxacillin-sensitive phenotype were removed to focus the analysis on MRSA, as oxacillin belongs to the same drug class it shares its mode of action with methicillin and has replaced methicillin in clinical use. 23 further isolates which possessed no phenotypic data and were predicted as MSSA during analyses were discarded. Additionally, 131 duplicate genomes were removed. Two CCoS isolates were removed for low read depth (<20x). Finally, 1207 MRSA genomes isolated in multiple Swiss hospitals and sequenced at the University Hospital of Basel between 2007 and 2022 were selected to represent a modern collection. After this process 451 MRSA isolates from the CCoS, 103 from public repositories (18 from Pathogenwatch and 85 from NCBI Pathogen Detection), and 1207 from the USB were used in further analysis.

### Genotyping

This work used two genotypic typing methods: multi-locus sequence typing (MLST) and core genome MLST (cgMLST, (38)). The genomes were typed in Ridom SeqSphere+ (v8.3.0), generating MSTs of MLST or cgMLST data, and a world map displaying the origin of the isolates (Map data © Google, INEGI). For all visualizations, colorblind-friendly color palettes were created with the “coolors” website (80). cgMLST clusters were generated with a minimum cluster size of 15 and a maximum cluster distance of 24 different alleles (38).

### Phylogenies

To improve visualization comprehensibility for this step, the 1207 genomes from the USB were dereplicated with a 0.005 threshold in Assembly Dereplicator (v0.1.0) (81) resulting in 223 genomes. Assembly dereplicator clusters the genomes with dissimilarity lower than the threshold and keeps the assembly with the largest N50 for each cluster. An *S. epidermis* reference genome (NZ_CP035288.1) was added as an outgroup for rooting purposes. Prokka (v1.14.5) (82) was used to annotate the genomes. From the annotated genomes, a the core genome alignment was generated with roary (v3.13.0) (82).This core genome alignment was processed with IQ-TREE 2 (v 2.2.1) (83). Whole genome alignments of cgMLST clusters were calculated with SKA (v 1.0) (84), within-house hybrid assemblies used as a reference where available. For ST22, reference sequence NZ_CP053101.1 was used. From these alignments, phylogenies without recombination were generated with gubbins (v 3.2.1) (85) and rooted with BactDating (v1.1.1) (86). Clusters were investigated with fastBAPS (v 1.0.8) (87). The phylogenies were visualized in RStudio (v2022.07.1+554).

### AMR and virulence prediction

In order to predict AMR and virulence, parts of the-finder software suite developed by the Danish center for genomic epidemiology were used. The Resfinder Software (88) was applied to all genomes. Leveraging the antibiograms of the isolates, they were compared with the software prediction to gauge software performance. Virulencefinder (89) was also run on all genomes. The detected genes were arranged in a presence absence matrix. Subsequently the virulence genes were classified by using the virulence factors database VFDB (90), Uniprot (91) and by consulting relevant publications.

### SCC*mec* cassettes

SCCmecfinder was also implemented for the genome set after dereplication using the SCCmecfinder web server (92).

### R packages

The following R packages were used: pacman (v0.5.1) (93), tidyverse(v1.3.4) (94), rjson (v0.2.21) (95), ggtree (v3.2.1) (96), treeio (v1.18.1) (97), tidytree (v0.4.1) (98), ape (v5.6.2) (99), flextable (v0.8.2) (100), BactDating (v1.1.1) (86), fastbaps (1.0.8) (87), svglite (v2.1.0) (101), knitr (v1.4.0) (102).

## Acknowledgments

We thank Elisabeth Schulthess, Christine Kissling, Magdalena Schneider, Valerie Courtet, Daniel Gandner, Sabrina Stammler, Anne Schnäpf, Selda Turkan, Brigitte Friedli, and Rosamarie Vesco for their excellent technical assistant in sequencing the MRSA isolates and maintaining the isolate collection (all USB). We also thank Prof. Georg Lipps (Applied School of Science, Muttenz) for scientific inputs during the Bachelor thesis of Vanni Benvenga. We also thank Prof. em. Kayser, who donated this valuable isolate collection to the CCoS. Calculations were performed at sciCORE (http://scicore.unibas.ch/) scientific computing center at University of Basel.

## Glossary

MRSA: methicillin resistant *Staphylococcus aureus*
MSSA: methicillin sensitive *Staphylococcus aureus*
HA-MRSA: hospital-associated
MRSA CA-MRSA: community-associated
MRSA LA-MRSA: livestock-associated
MRSA CCoS: Culture Collection of Switzerland
USB: University Hospital Basel
MLST: multilocus sequence type
cgMLST: core genome multilocus sequence type
ST: sequence type
CC: clonal complex
SLV: single locus variant
CI: confidence interval

